# Dendrites decrease the synaptic weight resolution necessary to implement linearly separable computations

**DOI:** 10.1101/2020.04.20.051342

**Authors:** Romain Cazé, Marcel Stimberg

## Abstract

In theory, neurons modelled as single layer perceptrons can implement all linearly separable computations. In practice, however, these computations may require arbitrarily precise synaptic weights. This is a strong constraint since both, biological neurons and their artificial counterparts, have to cope with limited precision. Here, we explore how the non-linear processing in dendrites helps overcoming this constraint. We start by finding a class of computations which requires increasing precision with the number of inputs in a perceptron and show that it can be implemented without this constraint in a neuron with sub-linear subunits. Then, we complement this analytical study by a simulation of a biophysical neuron model with two passive dendrites and a soma, and show that it can implement this computation. This works demonstrates a new role of dendrites in neural computation: by distributing the computation across independent subunits, the same computation can be performed more efficiently with less precise tuning of the synaptic weights. We hope that this works not only offers new insight into the importance of dendrites for biological neurons, but also paves the way for new, more efficient architectures of artificial neuromorphic chips.

**Author Summary:** In theory, we know how much neurons can compute, in practice, the number of possible synaptic weights values limits their computation capacity. Such a limitation holds true for artificial and synthetic neurons. We introduce here a computation where the required means evolve significantly with the number of inputs, this poses a problem as neurons receive multiple thousands of inputs. We study here how the neurons’ receptive element-called dendrites-can mitigate such a problem. We show that, without dendrites, the largest synaptic weight need to be multiple orders of magnitude larger than the smallest to implement the computation. Yet a neuron with dendrites implements the same computation with constant synaptic weights whatever the number of inputs. This study paves the way for the use of dendritic neurons in a new generation of artificial neural network and neuromorphic chips with a considerably better cost-benefit balance.

## Introduction

In theoretical studies, scientist typically represent neurons as a linear threshold units (LTU; summing up the weighted inputs and comparing the sum to a threshold) [11]. Multiple decades ago, theoreticians exactly delimited the computational capacities of LTU also known as the Perceptron [12]. LTU cannot implement computations like the exclusive (the XOR), but they can implement all possible linearly separable computations and a sufficiently large network of LTUs can approximate all possible computations.

Research in computer science determined the synaptic wights resolution necessary and sufficient to compute all linearly separable computation [13, 8], and they evolves exponentially with the number of inputs. Formally an LTU needs integer-valued weights so that 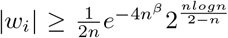, with *β* a constant, to be able to implement all linearly separable functions [8]. Consequently, an LTU with finite means cannot compute all linearly separable functions [5].

This limitation poses multiple practical problems. In the nervous system, neurons need to maintain a large number of synapses or synapses with a large number of stable state. Neuromorphic chips illustrate the problem and synapses often occupy the majority of the space, up to ten times more than the space occupied by neurons themselves [14]. We demonstrate here that dendrites might be a way to cope with this challenge.

Dendrites are the receptive element of neurons where most of the synapses lay. They turn neurons into a multiple layers network [15, 20] because of their non-linear properties [1, 16]. They enable neurons to compute linearly inseparable computation like the XOR or the feature binding problem (FBP) [6, 3]. We wonder here if dendrites can also decrease the synaptic resolution necessary to compute linearly separable computations.

First, we investigate the three inputs variables computations implementable by an LTU with positive synaptic weights. Second, we extend the definition of one of this computation to an arbitrarily high number of inputs. Third, we implement this computation at a smaller cost in a neuron with two passive dendrites.

This work not only shows the usefulness of dendrites in the nervous system, but also paves the way for a new generation of more cost-efficient artificial neural network and neuromorphic chips composed of dendritic neurons.

## Materials and methods

### Biophysical neuron model

We performed simulations in a spatially extended neuron model, consisting a spherical soma (diameter 10 μm) and two cylindrical dendrites (length 400 μm and diameter 0.4μm). The two dendrites are each divided into four compartments and connect to the soma at their extremity. In contrast to a point-neuron model, each compartment has a distinct membrane potential.

The membrane potential dynamics of the individual compartments follow the Hodgkin-Huxley formalism with:

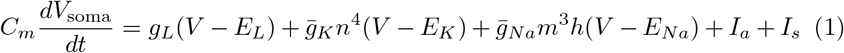

for the somatic compartment and

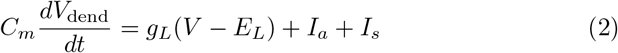

for the dendritic compartments.

Here, *V*_soma_ and *V*_dend_ are the respective membrane potentials, *C_m_* = 1 μF cm^-2^ is the membrane capacitance, 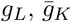, and 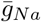 stand for the leak, the maximum potassium and sodium conductances respectively, and *E_L_, E_K_*, and *E_Na_* stand for the corresponding reversal potentials. The currents *I_a_* represent the axial currents due to the membrane potential difference between connected compartments. The synaptic current *I_s_* arises from a synapse placed at the respective compartment. It is described by

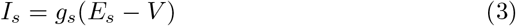

with *E_s_* being the synaptic reversal potential and *g_s_* the synaptic conductance. This conductance jumps up instantaneously for each incoming spike and decays exponentially with time constant *τ_s_* = 1 ms otherwise:

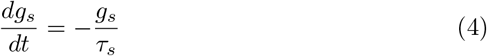

The dynamics of the gating variables *n, m,* and *h* are adapted from [19] and omitted here for brevity.

The parameter values are summarized in Table 1

**Table 1:**
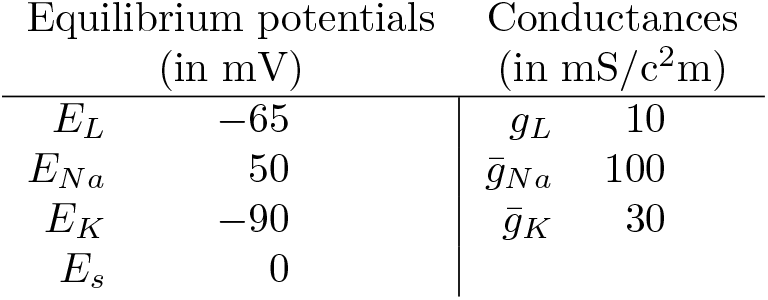
Parameter values used in the biophysical model

| Equilibrium potentials<br>(in mV) |  | Conductances<br>(in mS/c<sup>2</sup>m) |  |
| --- | --- | --- | --- |
| $E_L$ | -65 | $g_L$ | 10 |
| $E_{Na}$ | 50 | $\bar{g}_{Na}$ | 100 |
| $E_K$ | -90 | $\bar{g}_K$ | 30 |
| $E_s$ | 0 | | |

Note that due to the absence of sodium and potassium channels in the dendrites, the dendrites are passive and cannot generate action potentials.

All simulations were performed with Brian 2 [18]. The code is available as a supplementary zip file. It allows reproducing the results presented in Fig 4, Fig 5 and Fig 6. To demonstrate that the details of the neuron model do not matter for the results presented here, the code can also be run with a simpler leaky integrate-and-fire model.

### Elementary neuron model and Boolean functions

We first define as a reminder Boolean functions:

#### Definition 1.

*A Boolean function of n variables is a function on* {0, 1 }^*n*^ *into* {0,1}, *where n is a positive integer.*

Note that a Boolean function can also be seen as a binary classification or a computation.

A special class of Boolean functions which are of particular relevance for neuron are linearly separable computations:

#### Definition 2.

*f is a linearly separable computation of n variables if and only if there exists at least a vector w* ∈ ℝ^*n*^ *and a threshold* Θ ∈ ℝ *such that*:

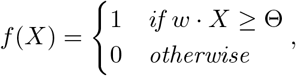

*where X* ∈ {0, 1}^*n*^ *is the vector notation for the Boolean input variables.*

Binary neurons are one of the simplest possible neuron models and closely related to the functions described above: their inputs are binary variables, representing the activity of their input pathways, and their output is a single binary variable, representing whether the neuron is active or not. The standard model is a linear threshold unit (LTU), defined as follows:

#### Definition 3.

*An LTU has a set of n weights* 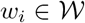 *and a threshold* 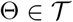 *so that*:

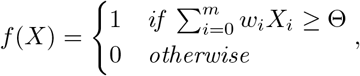

*where X* = (*X*_1_,…, *X_m_*) *are the binary inputs to the neuron, and* 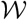 *and* 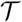 *are the possible values for synaptic weights and the threshold, respectively.*

This definition is virtually identical to Def. 2, however, *w***i** and Θ are no longer arbitrary real values, but chosen from a finite set of numbers depending on the implementation peculiarities and noise at which these value can be stabilised. It means that a neuron may not be able to implement all linearly separable functions. For instance, a neuron with non-negative weights can only compute positive linearly separable functions:

#### Definition 4.

*A threshold function f is positive if and only if f* (*X*) ≥ *f* (*Z*) ∀ (*X,Z*) ∈ {0, 1}^*n*^ *such that X ≥ Z* (*meaning that ∀i: x_i_ ≥ z_i_*)

To account for saturations occurring in dendrites, we introduce the sub-linear threshold unit (SLTU):

#### Definition 5.

*A SLTU with d dendrites and n inputs has a set of d × n weights W_i,j_* ∈ {0, 1} *with n w_i_ such that ∑_j_ w_i,j_* = *w_i_, d dendritic threshold* 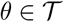 *and a threshold* 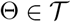, *such that*:

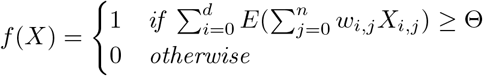

*with*

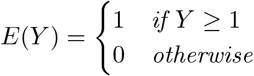

Such a neuron model can compute all positive Boolean functions (see Def. 4) provided a sufficient number of dendrites and synapses [3].

We used integer-valued and non-negative parameters both for the LTU and the SLTU without loss of generality. It allows to exactly determine the number of similar synapses necessary to implement a given computation.

## Results

### Function implementations for three input variables

In the following, we will look at computations that cannot be implemented in an LTU without using different strictly positive synaptic weights. The simplest case where this can occur is for three inputs. We list all such computation in Tab 2. Of these computations, the first three (OR, AND/OR, and AND) do not require distinct synaptic weights; they can all be implemented in an LTU by having the same weight for all synapses, by only varying the threshold. This is not the case for the last two computations that we coined the Dominant AND (D-AND) and the Dominant OR (D-OR): here, an LTU implementation needs to have one synaptic weight that is twice as big as the others (see Fig. 1).

**Figure 1:**
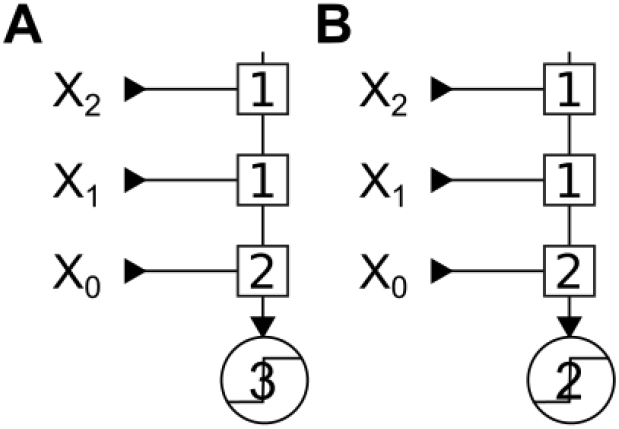
Minimal implementation of the Dominant AND computation (D-AND) and its dual by a linear threshold unit (LTU). Implementations of the D-AND where *X*_0_ is the dominant input. Squares represent synapses with their synaptic weight, and circles stand for transfer functions. Here, the transfer functions are threshold functions with the given value as their threshold. A: Implementation of the D-AND, note that *X*_0_ has twice the synaptic weight compare to the others. B: Implementation of the D-OR, note that we keep the same synaptic architecture and we only change the threshold of the transfer function.

**Table 2:** The five computations for *n* = 3 inputs with their associated truth tables. We have assigned a name to each class for easier reference.

| Inputs | OR | AND/OR | AND | D-OR | D-AND |
| --- | --- | --- | --- | --- | --- |
| 000 | 0 | 0 | 0 | 0 | 0 |
| 001 | 1 | 0 | 0 | 0 | 0 |
| 010 | 1 | 0 | 0 | 0 | 0 |
| 011 | 1 | 1 | 0 | 1 | 0 |
| 100 | 1 | 0 | 0 | 1 | 0 |
| 101 | 1 | 1 | 0 | 1 | 1 |
| 110 | 1 | 1 | 0 | 1 | 1 |
| 111 | 1 | 1 | 1 | 1 | 1 |

We have named the first new computation the D-AND because it requires the activation of a dominant (D) input AND the activation of another input. The D-OR is the Boolean dual of the D-AND, i.e. obtained by replacing AND operations by OR, and vice versa. In this function, an activation of the dominant input alone triggers an output. The other way to trigger an output is to activate both *X*_1_ and *X*_2_ at the same time. These functions have a “dominant input” – an input that is *sufficient* to make the output true (D-OR), respectively *necessary* to make the output true (D-AND).

In the present paper, we always chose *X*_0_ as the dominant input, but we could have picked *X*_1_ or *X*_2_. There is nothing comparable in the other three computations which treat all inputs identically.

An LTU (Fig. 1) implements the computation by using synaptic strength. We employed here integer valued synaptic weights to reflect their finite precision. Even if synaptic weights can take real values, a finite precision means a finite number of values, and one could consider each integer to represent the index of this value. The weight and threshold values to implement a function are obviously not unique. For example, we could multiply all the weights by 2 and set the threshold to 6 (D-AND), or 4 (D-OR) and obtain the same results. Here we always use the lowest possible integer valued synaptic weights, and the corresponding lowest possible threshold.

Then, we wanted to implement the D-AND and D-OR computation in threshold units with non-linear dendritic subunits, as an abstraction of neurons with dendrites [15].

We consider two types of non-linearities: a threshold function to model supra-linear summation; and a saturating function to model sub-linear summation (SLTU; see Material and Methods). Both types of summations have been observed in dendrites. Dendritic spikes are a well-known example of supra-linear summation [6], while sub-linear summation can be observed in completely passive dendrites due to the intrinsic saturation of synaptic conductances [1].

Fig. 2 shows a minimal implementation of D-AND in a dendritic neuron, with a supra-linear summation in Fig 2A and a sub-linear summation (SLTU) in Fig 2B. In both cases, all synapses are of identical strength. However, note that in the supra-linear implementation in Fig 2A, the *X*_0_ input connects to both dendrites. Therefore, as we define an input’s synaptic weight as the total effect it has in the final summation stage (analogous to depolarisation measured in the soma of a neuron), we have to consider the weight of *X*_0_ as twice as high as the other inputs. Note that this makes this implementation similar to the implementation in an LTU (Fig 1A): the dominance of *X*_0_ is expressed by a stronger weight. This is not the case for the sub-linear implementation in Fig 2B, where the synaptic weights are identical. Here, the dominance of *X*_0_ is only expressed by its placement.

**Figure 2:**
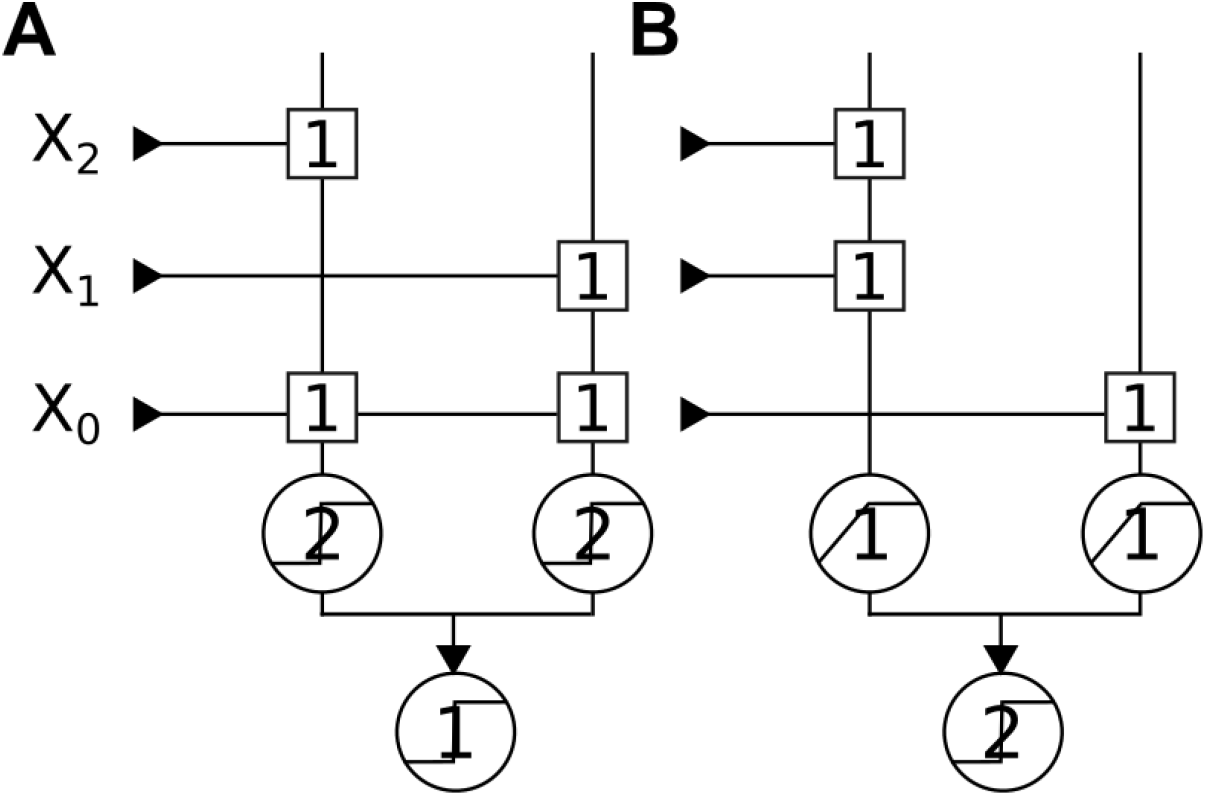
Two implementation of the D-AND in threshold units with non-linear sub-units (“dendritic neurons”). Squares represent synapses and circles represent transfer functions and their respective threshold/saturation values. Note that the final transfer functions (“somatic integration”) are always threshold units, whereas the transfer functions of the sub-units (“dendrites”) are threshold functions as an example for supra-linear summation in A, and saturating functions as examples of sub-linear summation in B. A: D-AND Implementation using supra-linear summation. Note that *X*_0_ has two synapses and therefore a synaptic weight of two. B: D-AND Implementation using sub-linear summation. Here, all inputs have identical synaptic weights.

Since the D-OR is the dual of the D-AND, its implementation also follows the same structure as the D-AND, but with the sub- and supra-linear implementations reversed. In this case, the supra-linear implementation uses placement to implement the dominance of *X*_0_ whereas the sub-linear implementation uses strength.

In real neurons, spiking and saturating dendrites both integrate inputs sub-linearly in a given range [1, 6]. Therefore, we will focus on D-AND in the following section and further explore the potential of threshold units with sublinear sub-units (SLTUs) as an abstraction of dendritic integration in neurons.

### Implementing the D-AND for an arbitrary number of input variables

In the previous section, we have limited our analysis to computations with three input variables. We will now extend the definition of the D-AND to an arbitrary number of input variables. As in the three-variable case, we will consider one input to be the dominant input (assumed to be *X*_0_, without loss of generality). This input has to be activated together with at least one of the non-dominant inputs. Formally, we therefore define *f_n_*(*X*) as follows:

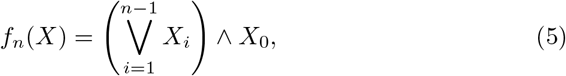

where *X* is the n-dimensional input vector with elements *X*_0_ … *X_n_*.

We can implement this function in an LTU (Fig 3A), as well as in a SLTU (Fig 3B)

**Figure 3:**
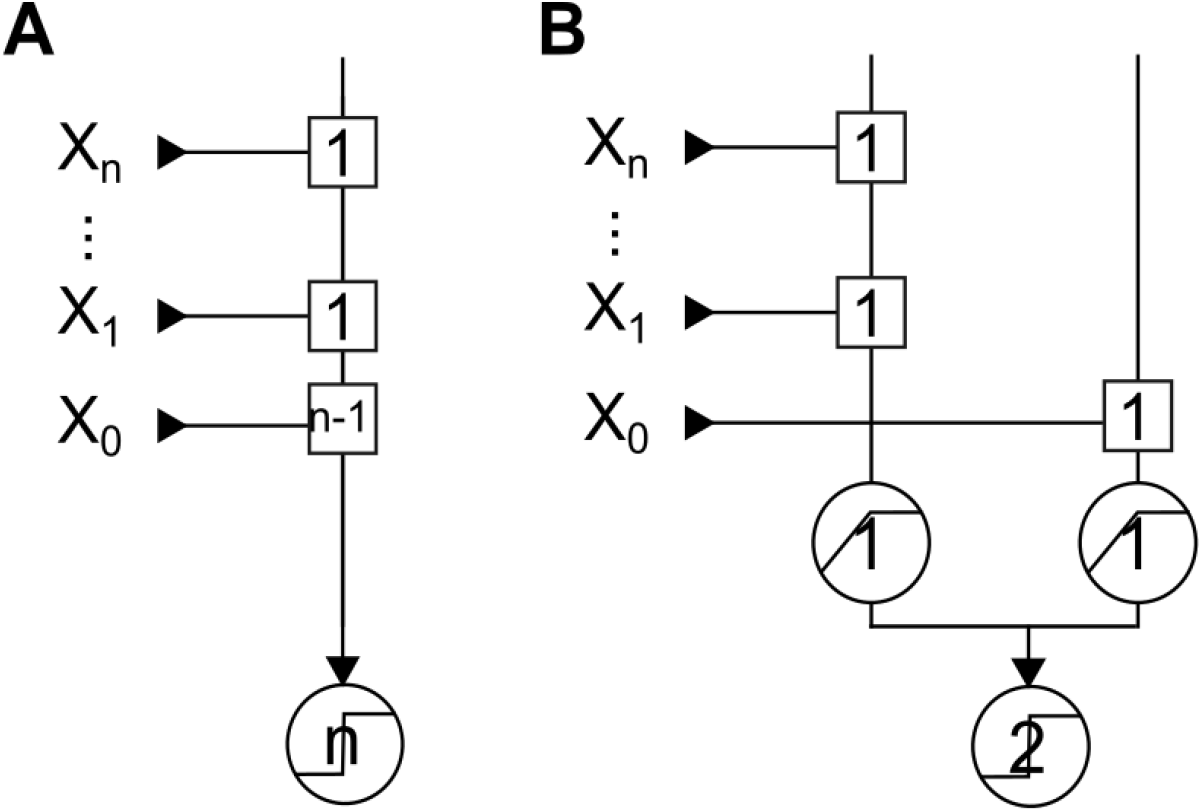
Extending the D-AND implementation to *n* inputs. Synaptic weights are in squares, and transfer functions are in circles. A: Minimal D-AND implementation in an LTU. Note that this implementation requires a synaptic weight that is *n* – 1 times bigger than the smallest weight. B: Implementation in an SLTU with sub-linear summation.

In the LTU implementation (Fig 3A), the D-AND of *n* variables requires that an input has a synaptic weight at least n — 1 times bigger than the other inputs, and the threshold has to grow accordingly.

We can summarise these observations in a proposition.

#### Proposition 1.

*To implement the D-AND, an LTU requires that an input has a synaptic weight n — 1 times bigger than the smallest synaptic weight.*

*Proof.* The LTU must stay silent when *X*_0_ is not active, even if *X*_1_, *X*_2_,…, *X*_*n*-1_ are active. Therefore *w*_1_ + *w*_2_ + … + *w*_*n*-1_ < Θ, thus Θ must be greater or equal than *n* × *w_m_* in with *w_m_* in the smallest synaptic weight.

Conversely, the output should be active as soon as *X*_0_ is co-active with any other input *X_j_* (for *j* ≥ 1). So *w*_0_ + *w_m_* in ≥ Θ, this means *w*_0_ + *w_min_* ≥ *n×w_min_*, thus *w*_0_ ≥ *w_min_*(*n* — 1).

In contrast, Fig 3B provides a constructive proof that an SLTU can implement the D-AND with equal synaptic weights. In this implementation, the distinguishing feature of the dominant input is that it targets the second dendrite; the synaptic weights and the threshold do not have to change with the number of inputs. If one would only measure the response to single inputs at the “soma” (last stage of summation), the dominant input would be indistinguishable from the other inputs, despite its dramatically different importance.

We will see next how these insights transfer to a more realistic biophysical model.

### Implementation of the D-AND in a biophysical model

We are going to use here an illustrative example: consider a predator of a certain shape (triggering input *X*_0_) and either coloured green or blue (inputs *X*_1_ and *X*_2_). For an animal that is a potential prey, a stimulus should only trigger a fleeing response if the predator’s shape is accompanied by one of the correct colours. Neither its shape alone nor either or both of the colours without the shape should trigger it.

Fig 4A presents a single neuron biophysical model implementing this function. One synapse comes from an input encoding for the shape of an object, and this synapse connects to the upper dendrite. The two others encode for two different colours, either green or blue, and they connect to the lower dendrite.

**Figure 4:**
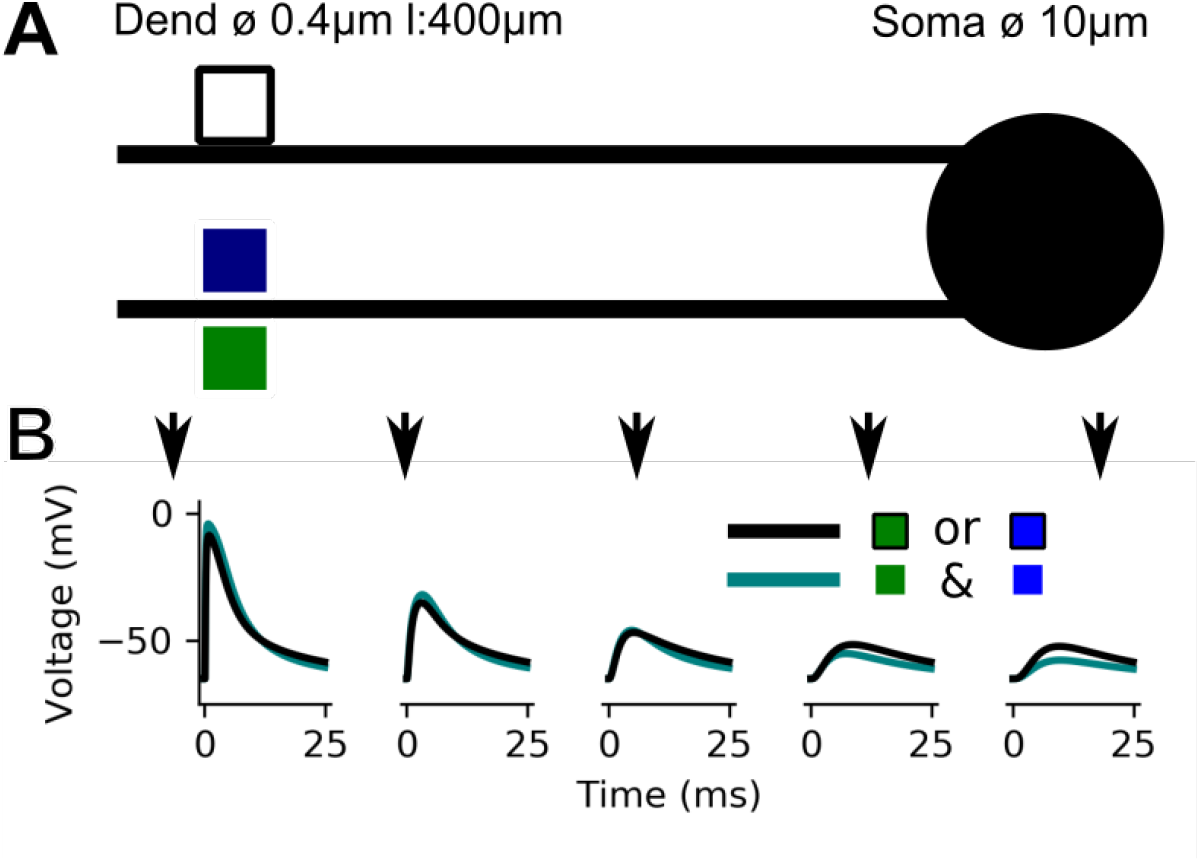
A biophysical model sensitive to synapses’ spatial distribution. A: A biophysical model with two dendrites and a soma (lines: dendrites, circle: soma). Coloured squares depicts synapses. The model has three equivalent synapses. One comes from an input encoding the shape (black square), and two situated on the other dendrite encode the colour (blue or green). B: Membrane voltage traces responding to either clustered (the two synapses activate on the same dendrite; aquamarine) or dispersed (the two synapses activate on distinct dendrites; black) synaptic activation at five distinct locations (dendrite where two synapses cluster at 350μm, 250μm, 150μm, 50μm, and soma). Note that at the point where two synapses cluster the clustered activation is larger than the dispersed activation, whereas in the soma the opposite is the case.

All the synapses, taken individually, produce the exact same depolarisation at the soma because we place them at the same distance (350 μm) and give them the same maximal conductance (20 nS).

We first look at the sub-threshold behaviour by disabling the sodium channels 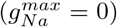. Fig 4B plots the voltage response at distinct locations on the lower dendrite, and finally in the soma. To probe the response we either activate a shape and one of the colours (black line Fig 4B), or both colours (aquamarine line). In both cases, we activate two synapses, but in the first case this activation is dispersed over the two dendrites whereas in the second case it is clustered on the lower dendrite. Despite activating the same number of synapses in both cases, and despite them all having the same strength, the depolarisation markedly differs. At the tip of the dendrite – where the synaptic stimulation takes place –, a dispersed activation leads to a smaller depolarisation than the clustered activation. In the soma, we observe the opposite: the dispersed ac-tivation leads to a bigger depolarisation than the clustered activation. For the dispersed activation, we record a depolarisation of 9.3 mV at the soma, whereas it is only 6.2 mV for the clustered activation.

We can explain this observation by considering the synaptic driving force [9]. The synaptic current induced by the activation of the synapse depends on the distance between the membrane potential and the synapses’ reversal potential; when several inputs drive the membrane potential closer to the reversal potential (here 0mV, this driving force diminishes. The combined effect of multiple synaptic inputs is therefore smaller then what is expected from summing the individual effects. In other words, the dendrite performs sub-linear summation.

Note that we could have expected a difference of up to 100% between the clustered and dispersed stimulation in the soma. In the actual simulation, the difference is closer to 50%, showing the non-trivial interaction taking place at the soma.

This means that even if all synapses have the same impact on the soma individually; even if we have a complete synaptic democracy [10], the relative placement of the synapses strongly influences the somatic response.

Based on the sub-threshold behaviour presented above, we will now show that we can implement the D-AND in a spiking neuron model. It is crucial to look at the supra-threshold behaviour as it is how the neuron communicate with the rest of the network. Moreover, back propagated action potentials might indeed undermine the dendritic non-linearity disrupting the implementation [2].

We can interpret Boolean inputs and outputs in different ways when we apply them to a biophysical spiking neuron model. Here, we will consider two interpretations. Firstly, we can think of an active input as corresponding to a continuous stimulation where the individual spikes arrive at random times, and of an active output as some spiking activity of the neuron (“rate interpretation”). Alternatively, we can think of active inputs as coincidentally arriving spikes within a certain time window, and accordingly of an active output as a single spike emitted in response (“spike interpretation”).

We present the model implementing the rate interpretation in Fig 5. We introduced this model earlier (Fig 4), except that it has now active sodium channels (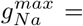; 650mScm^-2^). Each of its inputs (colours corresponding to the colours in Fig 4) activates 25 times according to a Poisson process.

**Figure 5:**
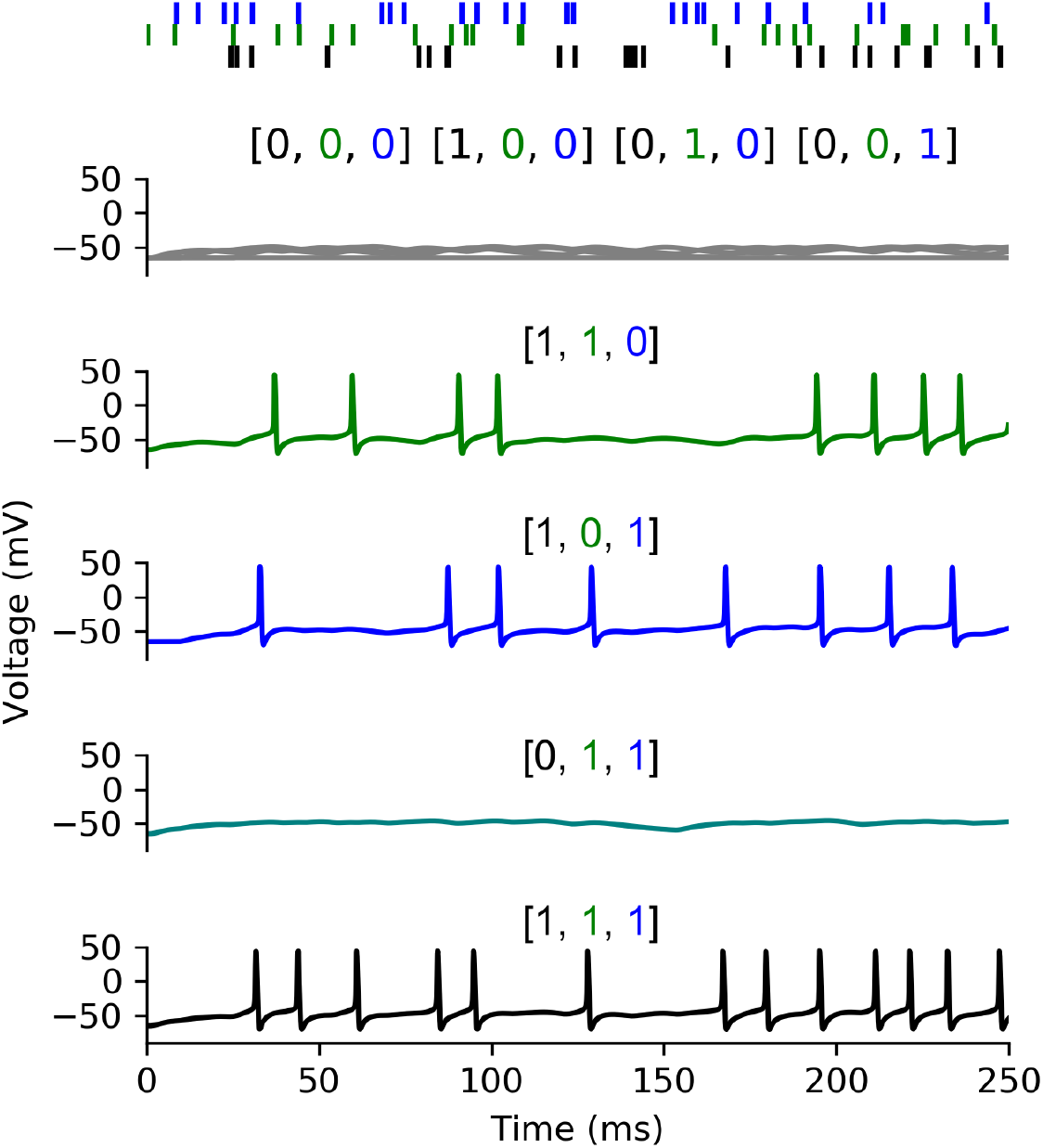
A biophysical model implementing the Dominant AND (rate interpretation) Top: activity of the three input synapses, the two first synapses impinge on the same dendrite while the black one impinges on another. Bottom: Eight somatic membrane responses depending on the active inputs. (gray: no synapse/only black/green/blue, green: black + green, blue: black + blue, aquamarine: green + blue, black: all inputs active). Note that this activity reproduces the truth table from Tab 2.

The Fig 5 displays, from top to bottom, the model’s responses in five different situations:

- A single input activates, in this case the neuron remains silent. We obtain the same outcome whatever the chosen input.
- Two dispersed inputs activate (black + green or black + blue), in these two scenarios the neuron fires.
- The two clustered inputs (green + blue) activate, in this case the neuron remains silent as expected from our observation in Fig 4B.
- All inputs activate, in this last case the neuron does not overly fire notably because of the refractory period.

This figure thus presents the response of the neuron model to all non-trivial cases, we have only omitted the case without any input activation (and therefore without any output activity).

Finally, we show an implementation of the spike interpretation in Fig 6. This model is identical to the model shown previously (Fig 5), except for a slightly lowered activation threshold of the sodium channels (*V_T_* = —55 mV instead of *V_T_* = —50 mV) to make it spike more easily. We discretize time into bins of 25 ms and decide randomly for each input whether it is active in each bin. If it is active, it activates at the beginning of the bin with a small temporal jitter (1 ms); inputs activating in the same bin therefore spike coincidentally. We can directly link these activations to Boolean variables that are either 0 (no spike) or 1 (spike). As Fig 6 shows, the neuron implements the D-AND and only spikes whenever the “shape input” (black) is active together with at least one of the “colour inputs” (blue and green).

**Figure 6:**
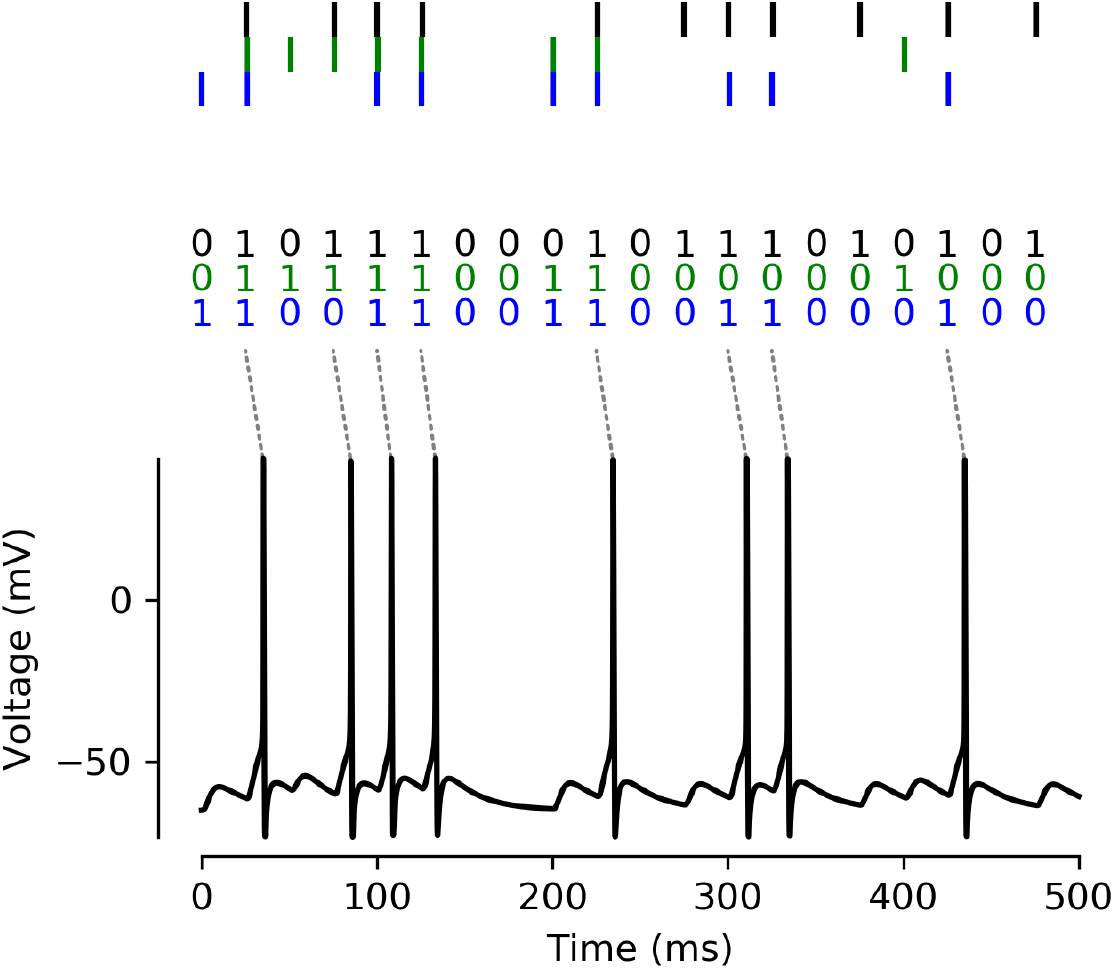
A biophysical model implementing the Dominant AND (spike interpretation) Top: The biophysical model receives input from three sources, where activation happens at regular intervals of 25 ms, with a random jitter of ±1 ms for each spike. We translate this activity into a binary pattern for each time bin of 25 ms. Bottom: The model’s membrane potential as measured in the soma. The response spikes implement the output of the D-AND computation as described in Tab 2.

We have shown that a biophysical model can implement the D-AND computation using a different strategy than the LTU. Each input has the same synaptic weight producing the same depolarisation at the soma. To distinguish between the inputs, the biophysical model uses location instead of strength: the dominant input (black) targets its own dendrite, while the two other inputs cluster on the same dendrite. With this strategy, the model can implement the D-AND. This implementation also works for two interpretations of the Boolean inputs and outputs — as elevated rates of spiking without temporal alignment,or as precisely timed coincident spikes.

## Discussion

In the present work, we demonstrate a computation for which an LTU implementation requires a synaptic weight many times higher than the other. In an implementation using dendrites, however, all synaptic weights can remain equal.

Two observations support this conclusion: 1) we have proven that an LTU requires that an input has a synaptic weight *n* — 1 times larger than the smallest input to implement a computation slightly more complex than coincidence detection. 2) A biophysical model with two dendrites can implement the same computation with all synaptic weights being equal.

The two first result sections look at a computation and one of its possible extension for an arbitrary number of uncorrelated inputs. Note that this extension of the D-AND from three to an arbitrary number of inputs is not the only possibility. We think that the extension we chose is a reasonable one and it allowed us to demonstrate an advantageous implementation in an SLTU compared to an LTU.

Note also that our denomination of one input as “dominant” and the others as “non-dominant” is very related to the distinction between “driver” and “modulator” inputs [17]. This concept, where driver inputs are necessary to activate a neuron, but this activity can be modulated by other inputs, is ubiquitous in the sensory system. For example, neurons in the primary visual cortex require a stimulus in their classical receptive field. Stimuli in the so-called extra-classical receptive field cannot activate the neuron by themselves, but strongly modulate the response if presented together with a stimulus in the classical receptive field [7]. This distinction is not entirely applicable in our example, since the dominant input *X*_0_ is not sufficient to activate the neuron by itself. Nevertheless, both computations rely on making a distinction between synaptic inputs, which can be implemented by placing inputs on different dendrites as we have shown in this study.

Our results also relate to a study [4] where we made use of dendrites to implement another computation with relevance to biology, namely direction selectivity. In the earlier study, we demonstrated how dendrites make the implementation resilient to massive synaptic failure, and this past work shows that dendrites improve the computing robustness. The present work introduces another aspect of robustness. Thanks to the saturating dendrite, the weights of the “non-dominant” synapses do not have to be set precisely; the only requirement is that a single input is enough to saturate the dendrite.

Several of our model properties fit with experimental observations. At least in some neurons, synapses at different positions tend to create the same depolarisation at the soma [10]. Furthermore, other experimental studies demonstrate examples of sublinear summation in dendrites [20, 1], notably in interneurons.

Our findings might also have implications beyond neuroscience, in particular for engineering applications. Studies in computer science assert that even problems solvable by an LTU might not have a solution when weights have a limited precision [5]. Being able to implement functions with equal synaptic weights, or with a small range of weights as we have demonstrated for the implementations in an SLTU, therefore may be advantageous for computations with limited resources.

In conclusion, dendrites enable to implement computations in more efficient and robust ways than without. This is not only of interest to neuroscientists, but might also inspire the design of future artificial neural networks and compact neuromorphic chips.

## Acknowledgements

This project started a long time ago within a team directed by Alain Destexhe it stems from multiple discussions with B. Teleńczuk. We also want to thank F. Danneville and C. Loyez for the stimulant discussions, Prof. A. Cappy for his remarks on the method, and M., Humphries for numerous valuable comments. Finally we want to thank A. Foust for making our scientific English easier to read and A. Marini-Audouard for the proof-reading before submission.

